# Fusion of quantitative susceptibility maps and T1-weighted images improve brain tissue contrast in primates

**DOI:** 10.1101/2021.10.05.462876

**Authors:** Rakshit Dadarwal, Michael Ortiz-Rios, Susann Boretius

## Abstract

Recent progress in quantitative susceptibility mapping (QSM) has enabled the accurate delineation of submillimeter scale subcortical brain structures in humans. The simultaneous visualization of cortical, subcortical, and white matter structure remains, however, challenging, utilizing QSM data solely. Here we present TQ-SILiCON, a fusion method that enhances the contrast of cortical and subcortical structures and provides an excellent white matter delineation by combining QSM and conventional T1-weighted (T1w) images. In this study, we first applied QSM in the macaque monkey to map iron-rich subcortical structures. Implementing the same QSM acquisition and analysis methods allowed a similar accurate delineation of subcortical structures in humans. However, the QSM contrast of white and cortical gray matter was not sufficient for an appropriate segmentation. Applying automatic brain tissue segmentation to TQ-SILiCON images of the macaque improved the classification of subcortical brain structures as compared to the single T1 contrast by maintaining a good white to cortical gray matter contrast. Furthermore, we validated our dual-contrast fusion approach in humans and similarly demonstrated improvements in automated segmentation of cortical and subcortical structures. We believe the proposed contrast will facilitate translational studies in nonhuman primates to investigate the pathophysiology of neurodegenerative diseases that affect subcortical structures such as the basal ganglia in humans.

**Highlights:**

- The subcortical gray matter areas of macaque monkeys are reliably mapped by QSM, much as they are in humans.
- Combining T1w and QSM images improves the visualization and segmentation of white matter, cortical and subcortical structures in the macaque monkey.
- The proposed dual contrast TQ-SILiCON provides a similar image quality also in humans.
- TQ-SILiCON facilitates comparative and translational neuroscience studies investigating subcortical structures.

## 1. Introduction

Precise delineation of brain structures plays an important role in neuroscience and medical imaging. Segmentation of brain structures is important for surgical planning and image-guided intervention and for the sequential monitoring of brain changes in health and disease. In particular, quantitative analyses of tissue morphology and clinical diagnosis strongly rely on the precise segmentation of cortical and subcortical structures. For example, during MRI-guided interventional studies in primates, brain tissue segmentation is a most valuable tool for the precise targeting and delivery of pharmacological and optogenetic agents into deep brain nuclei (Klink et al., 2021).

In both humans and nonhuman primates (NHPs), T1-weighted images (T1w) are widely used for the anatomical visualization of brain tissue structures. T1w provides excellent contrast for the segmentation of brain tissue classes such as gray matter, white matter, and cerebrospinal fluid. To better segment brain tissue, it is common to apply automatic volumetric and surface-based algorithms to the anatomical T1w datasets (Postelnicu et al., 2009). However, automated segmentation algorithms underperform on T1w images of NHPs, mainly due to inhomogeneities arising from the use of surface coils. Moreover, on T1w, subcortical structures appear very similar to white matter, hampering a correct segmentation. As a result, automatic tissue segmentation algorithms often misclassify subcortical deep gray matter nuclei as part of white matter tissue in humans and NHPs. Recent developments for segmentation of NHP T1w data focused on the use of deep-learning models (e.g. U-Net) (Wang et al., 2021), showing an improvement in brain segmentation over other methods implemented in the pre-processing pipelines of common analyses packages of FSL (Jenkinson et al., 2012), AFNI (Cox and Hyde, 1997) and Freesurfer (Fischl, 2012). Other approaches use no-contrast-based segmentation obtained from a nonlinear diffeomorphic averaging (Avants et al., 2011) of an NHP template that is used to nonlinearly warp individual NHP subjects along with their respective tissue types (Jung et al., 2021; Saleem et al., 2021; Saleem and Logothetis, 2012). This approach may, however, fail in case of significant deviations from the normal brain.

In contrast to T1w, Quantitative Susceptibility Mapping (QSM) provides vibrant contrast in subcortical deep gray matter nuclei, particularly within basal ganglia, due to the high iron abundance within these structures (Langkammer et al., 2012; Ramos et al., 2014). QSM is a rapidly evolving technique that uses gradient-recalled echo phase images to quantify the spatial distribution of magnetic susceptibility (Haacke et al., 2005, 2004). QSM contrast arises from the magnetic components in the tissue, such as iron, myelin, and calcium. The presence of myelin in the white matter results in diamagnetic susceptibility (Langkammer et al., 2012). In contrast, the presence of tissue iron in the form of ferritin macromolecules is the predominant contributor to the paramagnetic susceptibility of gray matter, including the cortex and subcortical structures. Significant progress in the post-processing of QSM data has enabled the accurate identification of small subcortical structures in humans (Guan et al., 2019; Schenck and Zimmerman, 2004). In particular, susceptibility changes around the basal ganglia have emerged as a potential biomarker for Parkinson’s disease and other neurodegenerative diseases (Guan et al., 2019; Schenck and Zimmerman, 2004; Shahmaei et al., 2019). However, our current understanding of the underlying sources that may give rise to contrast changes in basal ganglia and cortex remains elusive.

Given the challenges of identifying the mechanisms behind the contrast present in QSM of humans, developing QSM techniques in NHPs models is of particular value. NHPs have a brain organization similar to that of humans, and established knowledge exists about the anatomical and functional organization; for instance of the cortico-striatal-thalamic circuitry, much of which had been gained from immunohistochemistry (Hadaczek et al., 2016), pharmacological (Baron et al., 2002), neurophysiological and microsimulation (Nambu et al., 2015) techniques. Despite the gain in knowledge, the translation gap between non-invasive NHP research and human neuroimaging remains large. As an example, while QSM has been widely utilized in several human neuroimaging studies (Blezer et al., 2007; Haacke et al., 2009; Langkammer et al., 2016; Ravanfar et al., 2021; Shmueli et al., 2009), QSM has not been adopted as rapidly in the NHPs neuroimaging community (Dadarwal et al., 2019, 2021b; Yoshida et al., 2021).

In this study, we first applied QSM in the macaque monkey to map iron-rich subcortical structures. Given the rich contrast of QSM in subcortical structures and the gray-white matter contrasts available in T1w images, we aimed to develop a method for improving the visualization and segmentation of cortical and subcortical gray matter structures using the information provided by both T1w and QSM contrast. Here we focus on a contrast-based white and deep gray matter segmentation via the implementation of a fusion image, merging T1w and QSM by a linearly-weighted fusion algorithm. We term it T1w-QSM synthetic images via a linearly-weighted combination of contrasts (TQ-SILICON).

We believe our method has the potential to improve MRI-guided localization of deep brain stimulation sites in NHPs. To demonstrate the generalization of our approach, we implemented our dual-contrast acquisition in humans and showed similar improvements in brain tissue segmentation in both primate species.

## 2. Materials and Methods

### 2.1 Animal experiments

In total, 13 healthy, female, long-tailed cynomolgus macaques (*Macaca fascicularis*) have been used for this study. Four young adult monkeys at the age of 7.7 - 8.7 years (bodyweight 3.9 – 6.0 kg) were selected to define the respective weights for the linearly-weighted fusion of T1w and QSM. The obtained weights were then applied to 9 older macaques (age 16 - 18 years, bodyweight 4.3 - 8.1 kg) to validate the proposed synthetic tissue contrast.

All monkeys were purpose-bred, raised, and housed according to the standards for macaques at the German Primate Center (Göttingen, Germany). All aspects of the study were conducted in accordance with national and international guidelines of the German Animal Protection Law and the European Union Directive 2010/63/EU for the Protection of Animals used for Scientific Purposes. The study was approved by the local authorities, the Animal Welfare Service, Lower Saxony State Office for Consumer Protection and Food Safety (license-number 33.19-42502-04-16/2278).

In preparation for anesthesia, the macaques were deprived of food overnight. Anesthesia was induced by a mixture of ketamine (8.05 ± 2.65 mg per kg of body weight) and medetomidine (0.02 ± 0.01 mg per kg) and maintained by isoflurane (0.8 - 1.7% in oxygen and ambient air) via endotracheal tube and pressure-controlled active ventilation. The monkeys were placed in a prone position, and their heads were fixed in an MR-compatible stereotactic apparatus (Kopf 1430 M, https://kopfinstruments.com/product/model-1430m-mri-stereotaxic-instrument/).

### 2.2 Human volunteers

The proposed algorithm was applied to three male adult humans aged between 25 to 30 years. All measurements were performed after written informed consent. The human study was reviewed and approved by the ethics committee of the Georg-August-University of Göttingen.

### 2.3 MRI data acquisition

All data were acquired with a 3T MR system (MAGNETOM Prisma, Siemens Healthineers, Erlangen) equipped with a 7 cm single loop coil for macaque and a 20-channel head coil for human brain imaging. The imaging protocol included anatomical T1w and multi-echo gradient-recalled echo (ME-GRE) acquisitions. MR parameters for both macaques and humans are shown in **Table 1**.

**Table 1:** Macaque and human T1w and QSM data acquisition parameters. Abbreviations: TR - repetition time; TE - echo time.

| Macaque |  |  |
| --- | --- | --- |
| Parameters | T1w | QSM |
| Pulse sequence | 3D MPRAGE | 3D ME-GRE |
| Native resolution (mm<sup>3</sup>) | 0.5 x 0.5 x 0.5 | 0.31 x 0.31 x 0.31 |
| Field of view (mm<sup>2</sup>) | 108 x 128 | 97 x 120 |
| Matrix size | 216 x 256 | 312 x 384 |
| Total averages | 2 | 1 |
| Total acquisition time (min) | 14.3 | 24 |
| TR; TE (ms) | 2.7/2700 | 57/[3.7/4.9/48] |
| Inversion time (ms) | 850 | -- |
| Flip angle (degrees) | 8 | 20 |
| Pixel Bandwidth (Hz/Px) | 250 | 250 |
| **Human** |  |  |
| **Parameters** | **T1w** | **QSM** |
| Pulse sequence | 3D MPRAGE | 3D ME-GRE |
| Native resolution (mm<sup>3</sup>) | 0.8 x 0.8 x 0.8 | 0.75 x 0.75 x 0.75 |
| Field of view (mm<sup>2</sup>) | 300 x 320 | 195 x 240 |
| Matrix size | 240 x 256 | 260 x 320 |
| Total averages | 1 | 1 |
| Total acquisition time (min) | 6.3 | 7 |
| TR; TE (ms) | 2.2/2400 | 41/[4.5/4.5/36] |
| Inversion time (ms) | 1000 | -- |
| Flip angle (degrees) | 8 | 20 |
| Pixel Bandwidth (Hz/Px) | 220 | 260 |

### 2.4 MRI data analyses

T1w and ME-GRE images of both macaque and humans were converted to nifti format from dicom images using the function Dcm2niix (Li et al., 2016). ME-GRE magnitude images were pixel-wise averaged across echo times. The workflow for T1w and GRE analyses is shown in **Fig. 1A**. Single-subject T1w and mean ME-GRE magnitude images were denoised using ANTs *DenoiseImage* (Avants et al., 2011). The denoised volumes were then used to manually create brain masks using the ITK-SNAP segmentation tool (Yushkevich et al., 2006). We followed a manual segmentation process due to the unavailability of sufficient automatic whole-brain segmentation tools for the cynomolgus macaque brain. These brain masks were then used to correct T1w images from the bias fields using ANTs *N4BiasFieldCorrection* (Avants et al., 2011). The skull-stripped mean ME-GRE magnitude images were affinely registered (12 DOF) to the subject T1w brain volume using the ANTs registration algorithm.

**Figure 1:**
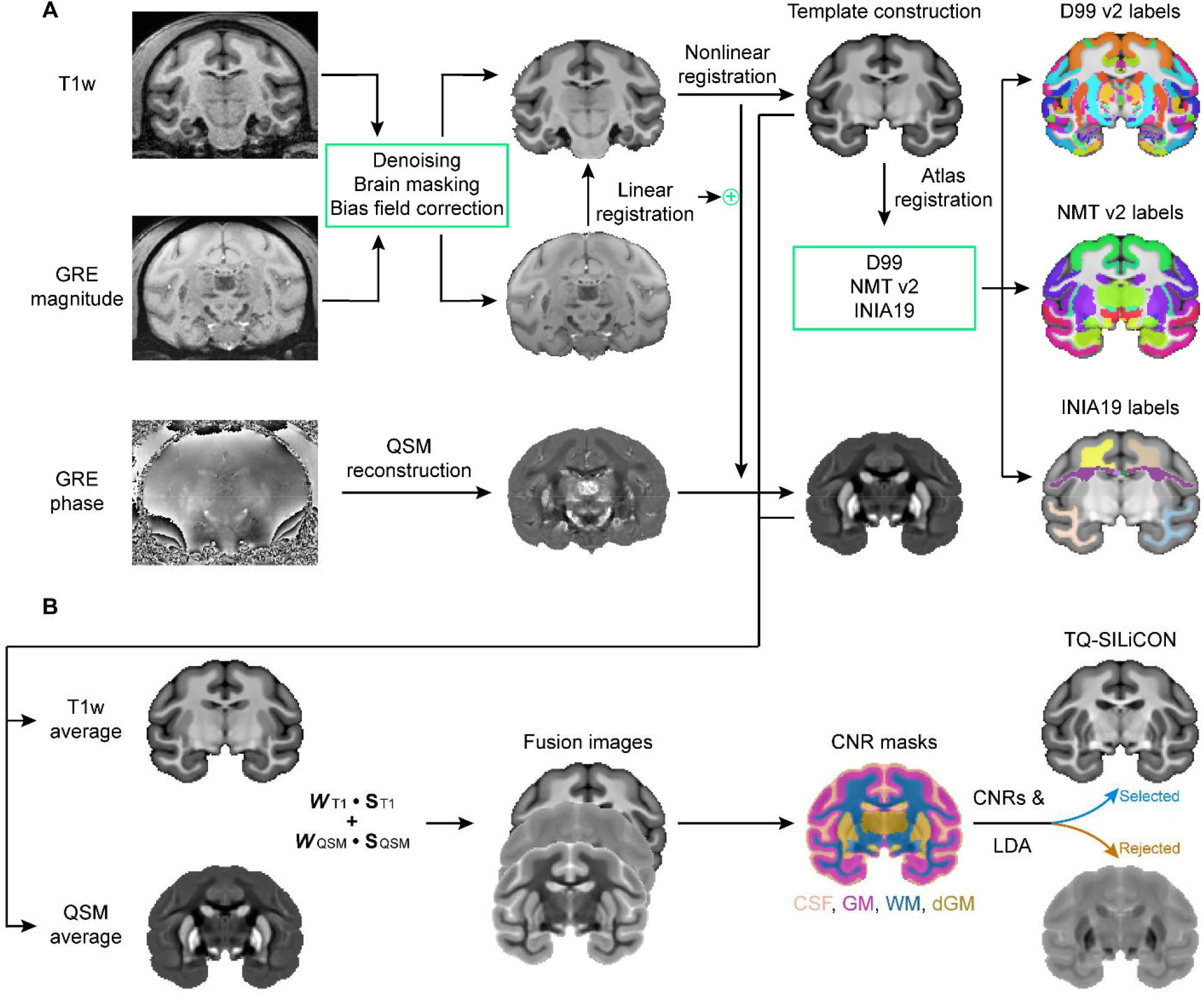
**A**. Analytical workflow for anatomical (T1w) and gradient-recalled echo (GRE) data. T1w and GRE magnitude images were denoised, brain masked, and corrected for field bias. QSM maps were reconstructed using the GRE phase images. The GRE magnitude brain image of the subject was linearly registered to the T1w image of the subject. All subject’s T1w images were registered nonlinearly to create a T1w average image, which was registered nonlinearly to the openly available macaque atlases (D99, NMT.v2, and INIA19). The atlases were used to extract cortical, subcortical, and white matter labels in T1w-average space. Subject GRE registration to T1w produced a linear registration transform, and subject T1w to T1w-average produced a nonlinear warping alignment. The combined transformations were then used to create an average QSM map. **B**. The TQ-SILiCON image generation workflow. T1w average and QSM average images were fused using a linearly-weighted combination. The respective weights (e.g., *WT1 and WQSM*) were randomly generated for the T1w and QSM images, while *ST1* and *SQSM* were then normalized by the signal intensities and susceptibility values of the T1w and QSM images, respectively. The generated fused images were first evaluated based on their contrast-to-noise ratio (CNR) between four tissue type classes (CSF, GM, WM, and deep GM). Images with sufficient CNR and with the highest linear discriminant analysis (LDA) accuracy score (e.g., classification to CSF, GM, WM) were then selected for further analysis. Abbreviations: GRE - gradient echo, CSF - cerebrospinal fluid, GM - gray matter, WM - white matter, and dGM - subcortical deep gray matter.

QSM maps of monkeys and humans were reconstructed using coil combined ME-GRE phase data. The QSM reconstruction included phase unwrapping using the best-path algorithm, background field removal using Laplacian boundary value and variable spherical mean value filtering algorithms, and solving the inversion problem using the multiscale dipole inversion approach (Abdul-Rahman et al., 2007; Acosta-Cabronero et al., 2018; Zhou et al., 2014).

T1w and QSM images from four macaques and three human subjects were duplicated and mirrored to the hemispheric plane, respectively. Using ANTs nonlinear registration (Avants et al., 2011), we constructed a symmetric population-averaged T1w template for both monkeys and humans by aligning all of the native and mirrored scans. The output deformation maps from the subject mean ME-GRE magnitude aligned to the symmetric population-averaged T1w template were then used to create symmetric population-averaged QSM templates. The macaque brain population-averaged T1w template was nonlinearly registered to the D99 (Saleem et al., 2021; Saleem and Logothetis, 2012), the NMT v2 (Jung et al., 2021), and the INIA19 (Rohlfing et al., 2012) macaque atlases.

A weighted linear combination was used to fuse symmetric population-averaged T1w and QSM templates, as shown in **Fig 1B**. The weights (*W*) for T1w were generated at random between 0 and 1, while weights for QSM were generated between −1 and 0. T1w signal intensities and QSM values (*X*) were normalized to a range between 0 to 1 before entering it into the fusion equation,

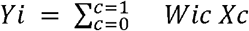

with *c* indicating the respective contrast (T1w or QSM) and *i* the number of iteration. In order to select the image fusion (*Y*), we used weights, the CNRs, and the linear discriminant analysis (LDA) accuracy score. Therefore, the three tissue classes (CSF, gray matter, and white matter) were derived from the labels of the available macaque atlases (D99 and INIA19). Where necessary, these labels were manually corrected and used to create respective tissue binary regions-of-interest (ROIs) masks using the ITK-SNAP tool (Yushkevich et al., 2006). Based on those tissue masks, the CNRs were estimated using the following formula:

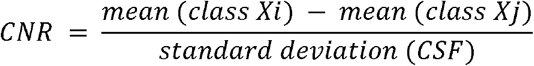

Where class *Xi* and *Xj* refer to the contrasts considered to estimate the CNR out of white matter, cortical gray matter, and subcortical gray matter. Using the same tissue masks, the LDA model was trained using *Scikit-learn* (Pedregosa et al., 2011) to identify the three tissue classes. We trained the model on 75% of the voxels and tested it on the remaining 25% of voxels. For the template brain, the minimum threshold for the CNR between gray matter and white matter was set to 3, while the threshold for the CNR between subcortical gray matter and white matter was set to 2. To maintain a lower contrast between the gray matter and the subcortical gray matter, the maximum CNR threshold for those was set to 0.5. Moreover, the threshold for the LDA accuracy score was set to 0.80. If the finally selected fusion image met all of the CNR requirements and obtained the highest LDA accuracy score out of 50 iterations, the image was selected as the TQ-SILiCON.

### 2.5 TQ-SILiCON evaluation

In order to compare TQ-SILiCON with T1w and QSM images, we calculated the CNR between (i) the motor cortex and the frontal white matter, (ii) the globus pallidus and the internal capsule, and (iii) the dentate nucleus and the cerebellum white matter. One-way ANOVA was performed in R to test for CNR differences between the three contrasts (e.g., T1w, QSM, and TQ-SILiCON). In case of significance, the CNR of TQ-SILiCON was compared with the CNR of T1w and QSM, respectively, using a two-tailed t-test. Significance levels were adjusted using the Bonferroni correction for multiple testing. To further evaluate the potential of the created fusion image for automatic tissue segmentation, ANTs Atropos was applied on single-subject T1w images and TQ-SILiCON of both macaques and humans. CSF was extracted beforehand using T1w, and a two-class segmentation (gray and white matter) was performed.

## 3. Results

This work aimed to establish multi-contrast imaging based on T1w and high-resolution QSM in NHPs. Below we describe our T1w and QSM results for identifying specific tissue structures in the cortex and subcortical gray matter nuclei. From the T1w and QSM images, we developed a fusion technique called TQ-SILiCON, enabling the improved automatic segmentation of white matter and cortical and subcortical gray matter. Importantly, we also evaluated our fusion strategy in humans and showed the potential of our approach for translational studies.

### 3.1 T1w and QSM in NHPs

T1w images of the macaque brain provided excellent gray-to-white matter contrast along the cortical surface. However, major subcortical deep gray matter nuclei remained challenging to delineate on T1w. In subcortical structures, T1w contrast largely delineated the caudate and putamen from the adjacent white matter, while the remaining deep gray matter nuclei – contrast-wise – were identical to the neighboring white matter (**Fig. 2**). In contrast, QSM substantially enhanced the visibility of subcortical deep gray matter nuclei from other parts of the macaque brain (**Fig. 2**). Enhanced contrast on the QSM map clearly delineated subcortical structures such as caudate, putamen, external and internal segments of globus pallidus, thalamus, substantia nigra, red nucleus, and dentate nucleus. Due to their paramagnetic contrast, all of these deep gray matter nuclei appeared bright on the QSM map compared to the surrounding dark appearing white matter areas with diamagnetic susceptibility (**Fig 2**).

**Figure 2:**
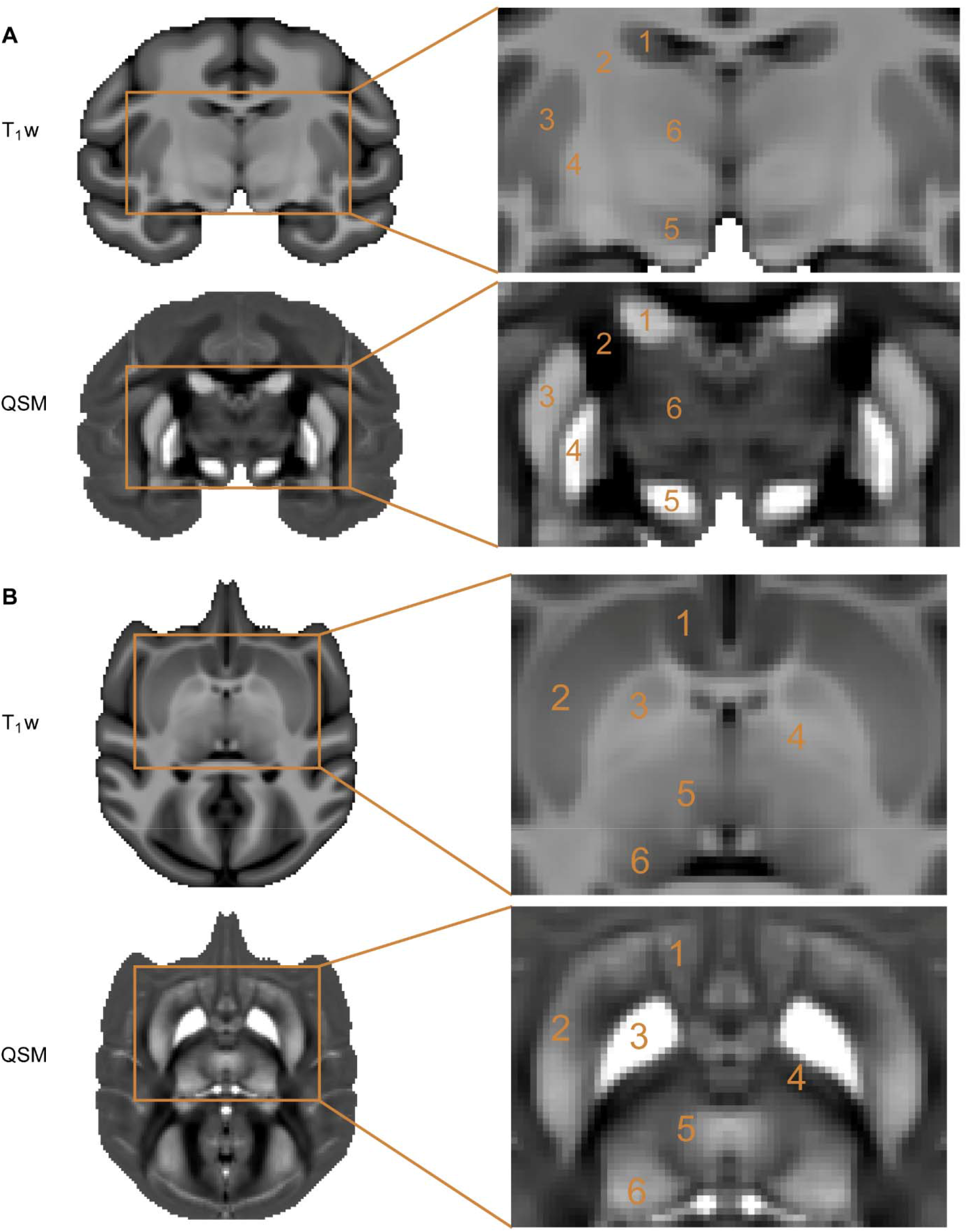
**A.** Complementary contrast between T1w and QSM images of the macaque brain. T1w images obtained from an average of 4 monkeys provide excellent contrast between white and gray matter. However, a magnified visualization of subcortical structures reveals relatively low contrast. On the other hand, QSM coronal images obtained from the average of the same 4 monkeys provide a unique contrast in subcortical structures but low contrast in the white and cortical gray matter. The locations of iron-rich subcortical structures are indicated on the magnified visualization of T1w and QSM images: 1 - caudate, 2 - internal capsule, 3 - putamen, 4 - globus pallidus, 5 - substantia nigra, and 6 - thalamus. **B.** T1w and QSM axial images of the macaque brain (average of 4 monkeys). The marked areas on the magnified visualization of T1w and QSM images are 1 - accumbens, 2 - putamen, 3 - globus pallidus, 4 - internal capsule, 5 - thalamus, and 6 - pulvinar.

Interestingly, the QSM contrast of the macaque thalamus highlighted different subnuclei, such as pulvinar, based on their varying susceptibility patterns (**Fig. 2B**). Among all subcortical deep gray matter nuclei in the macaque brain, the globus pallidus had the highest QSM contrast, followed by the substantia nigra, caudate, putamen, red nucleus, and thalamus, reflecting most likely a distinct amount of iron concentration. An important feature to highlight from the QSM contrast was the clear distinction between the internal and external segments of the globus pallidus.

While the QSM map provided improved visualization of the iron-rich subcortical deep gray matter (**Fig. 2**), QSM contrast lacked the high gray-white matter contrast as compared to T1w along the cortical surface and in deep brain areas such as those of the hippocampus. To circumvent the constraint of a single contrast, we pursued an analytical strategy that combines two MRI contrasts that are sensitive to different tissue substrates. Next, we demonstrate the linearly-weighted combination of T1w and QSM images which provides promising results applied to both NHPs and human neuroimaging data.

### 3.3 Merging T1w and QSM provides a 3D data set with enough contrast for segmenting subcortical and cortical tissue in NHPs

We devised a workflow for fusing T1w and QSM images (**Fig. 1B**), which uses a linearly-weighted combination of the two image contrasts. The data-driven approach selected a fusion image with weight combinations of 0.83 and −0.87 for the T1w and QSM images, respectively. Example images for other weights are shown in **Fig. S1.** The selected TQ-SILiCON image derived from the four-monkey-average brain had a CNR value of 3.3 between gray matter and white matter, 2.8 between subcortical gray matter and white matter, and 0.4 between gray matter and subcortical gray matter and an LDA accuracy score of 0.86.

In macaques, the average TQ-SILiCON showed both excellent gray-to-white matter and deep gray matter contrast. The fusion technique amplified the contrast in subcortical structures and maintained white matter landmarks in delineating subcortical structures from adjacent white matter tissue. In comparison to the average T1w image, TQ-SILiCON was superior in delineating subcortical structures (**Fig. 3A**). Anatomically, we were able to identify the following nuclei: Caudate, putamen, external and internal globus pallidus, thalamus, substantia nigra, red nucleus, and dentate nucleus. When comparing the average TQ-SILiCON image to the average QSM map, the TQ-SILiCON image showed exceptional gray-white contrast evidently not present in the QSM map (**Fig. 3B**).

**Figure 3:**
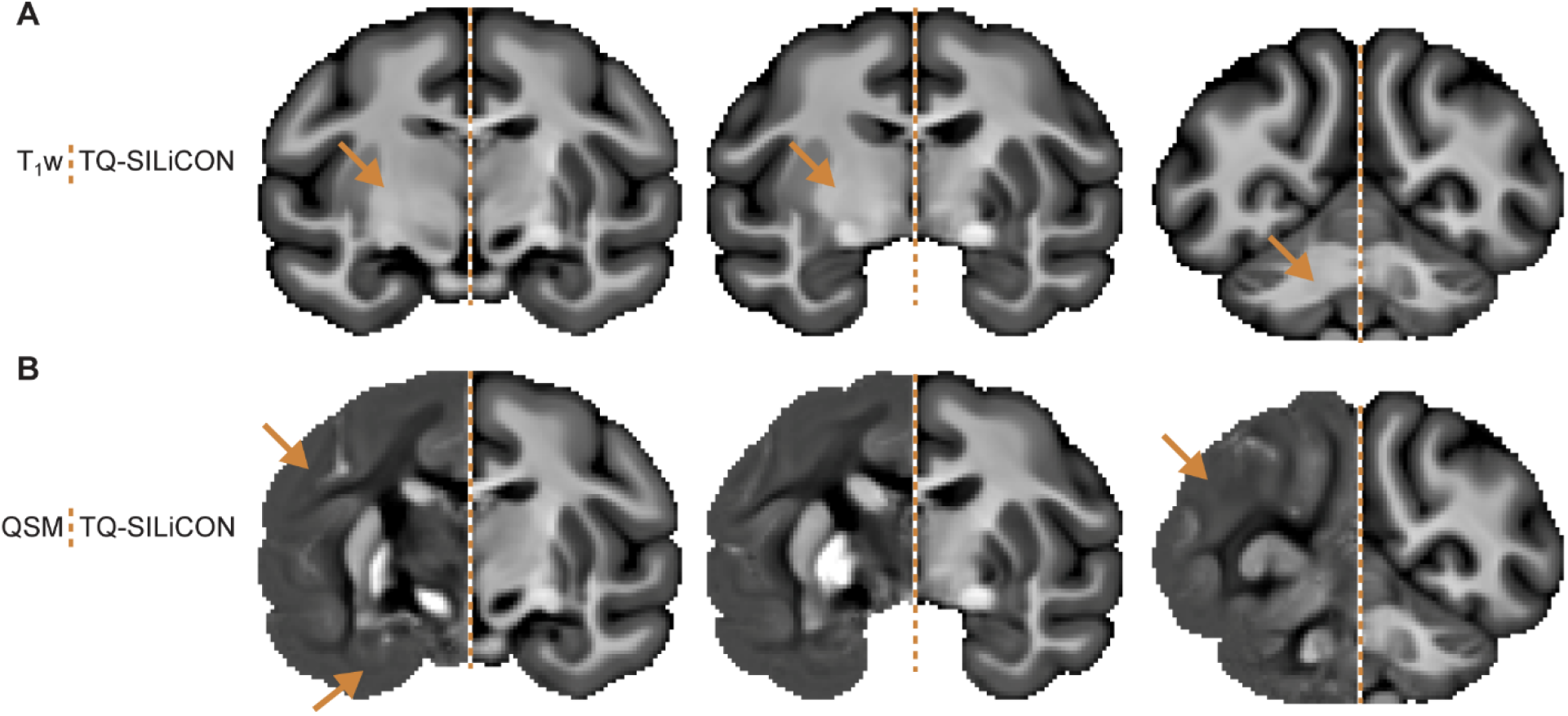
Enhanced cortical and subcortical contrast from TQ-SILiCON in the macaque monkey brain. **A**. Coronally oriented TQ-SILiCON (right) slices of the average macaque template as compared to the average T1w (left). **B**. Similarly, TQ-SILiCON (right) is compared to QSM images (left). In comparison to T1w, TQ-SILiCON has enhanced subcortical contrast while maintaining high gray and white matter contrast, which the QSM map lacks. Arrows point to areas in either T1w or QSM images that lack contrast.

To further demonstrate the applicability of our fusion technique to single-subject macaque datasets, we calculated TQ-SILiCON for all 13 monkeys separately using identical weights for all of them (**Fig. S7**). Compared to T1w, TQ-SILiCON revealed a significantly higher CNR between deep GM structures (globus pallidus p < 1e^−07^, dentate nucleus p < 1.1e^−06^) and the surrounding WM structure. In contrast, the CNR of the motor cortex and WM was similarly high for TQ-SILiCON and T1w and significantly higher as compared to QSM (p < 1e^−07^) (**Fig. 4C**). Thus, our quantification analyses revealed excellent gray-to-white matter contrast, including deep GM contrast at the single-subject level.

**Figure 4:**
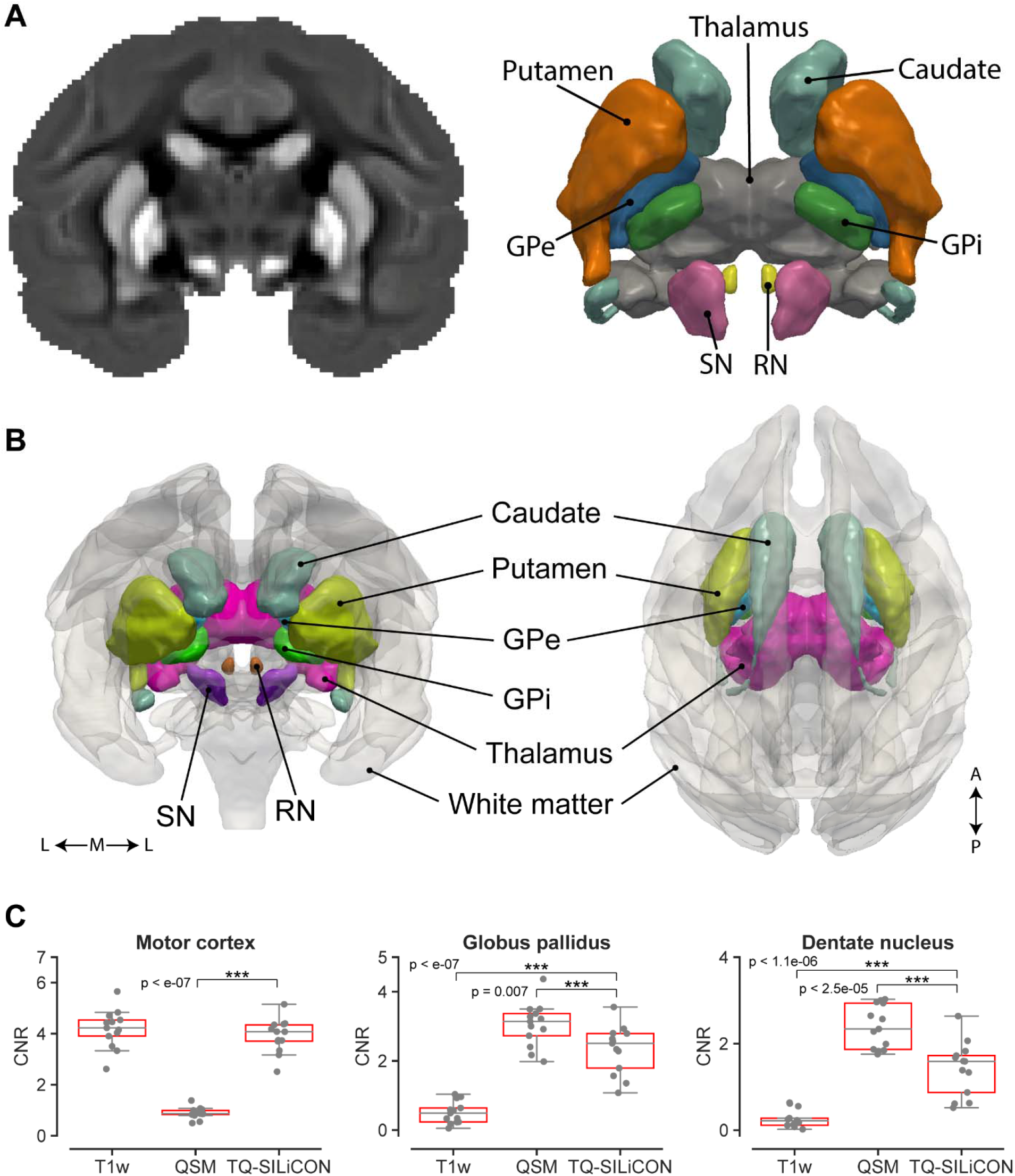
**A**. Improvement in cortical and subcortical segmentation after image fusion. **A**. QSM coronal image of the macaque brain showing the iron-rich subcortical structures on a 3D rendering: putamen, caudate, globus pallidus external (GPe), globus pallidus internal (GPi), thalamus, substantia nigra (SN), and red nucleus (RN). **B**. TQ-SILiCON allows whole-brain cortical and subcortical segmentation of the brain tissue. The white matter surface (transparent) of the macaque brain allows visualization of the segmented subcortical deep gray matter nuclei (colored) within the brain. Lateral (L) - Medial (M) - Lateral (L) and Anterior (A) - Posterior (P). **C**. The contrast to noise ratio (CNR) was calculated between the motor cortex and frontal white matter, the globus pallidus and internal capsule, and the dentate nucleus and cerebellum white matter. The CNRs were calculated using the absolute difference between the signal mean of the gray and white matter structures divided by the standard deviation of the respective white matter voxels. Level of significance p < 0.025.

We then explored how the enhanced CNRs may further improve the automatic tissue segmentation for gray matter and white matter for each contrast (T1w and TQ-SILiCON) on all the macaque datasets (**Fig. 5 and 6**). The segmentation based on TQ-SILiCON showed an improved gray-white matter delineation compared to those of T1w. T1w image-based segmentation misclassified some of the subcortical structures as white matter **(Fig. 5)** and some white matter regions as gray matter, including the extreme capsule, external capsule, and lateral medullary lamina, which were all correctly classified as white matter by TQ-SILiCON-based segmentation (**Fig. S2**). These findings were partially confirmed by calculating the number of misclassified voxels in each macaque dataset (**Fig. 6**). Whereas the number of WM voxels misclassification as GM was almost comparable between TQ-SILiCON and T1w, TQ-SILiCON produced significantly fewer misclassified voxels in the basal ganglia.

**Figure 5:**
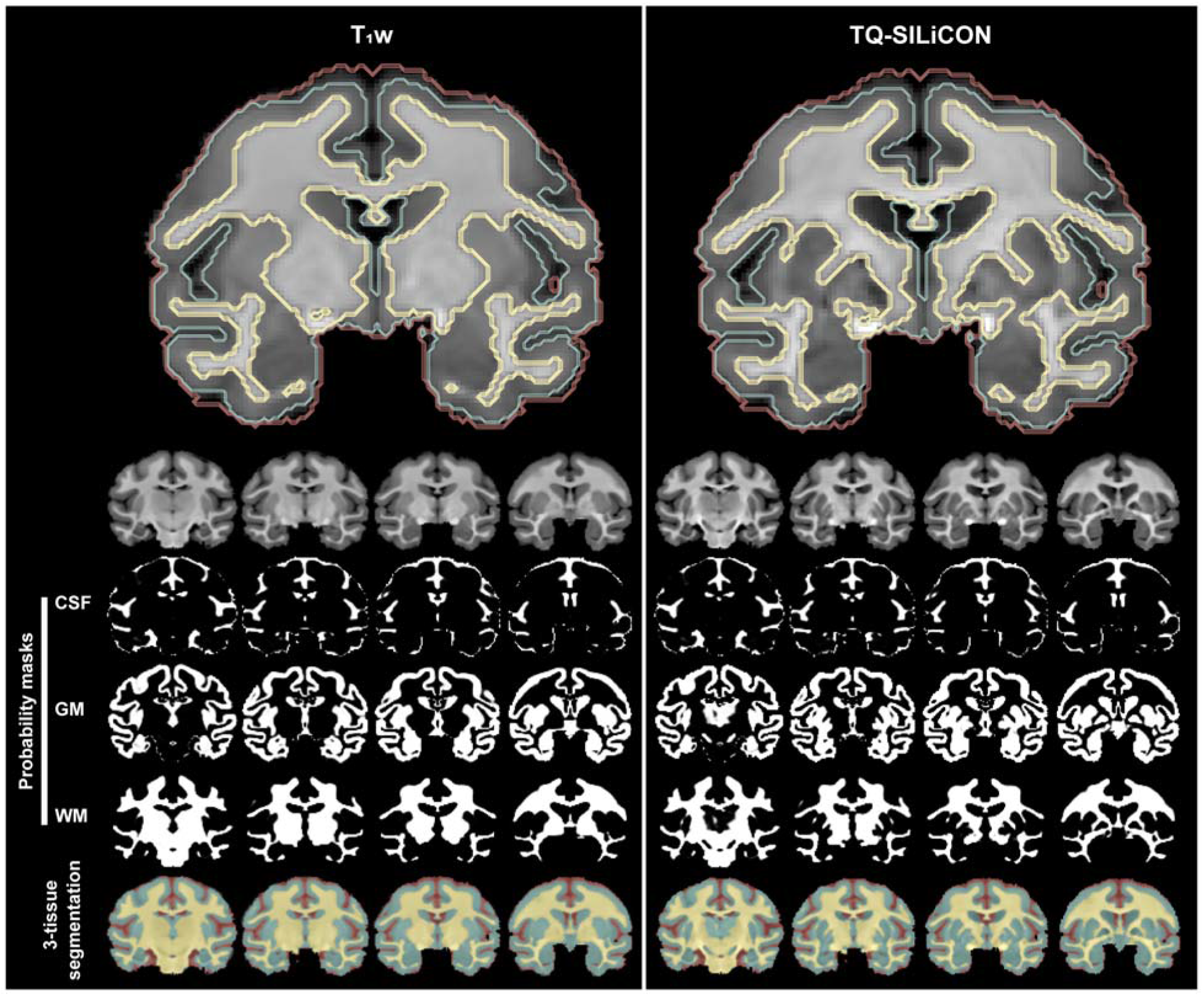
Improved automatic tissue segmentation after image fusion. (Left) Coronal view of a single macaque brain tissue segmentation using T1w and (right) TQ-SILiCON images. TQ-SILiCON enables better tissue classification of GM and WM. The bottom panel shows the semi-transparent color-coded tissue classification of the CSF, GM, and WM. Abbreviations: CSF - cerebrospinal fluid, GM - gray matter, and WM - white matter.

**Figure 6:**
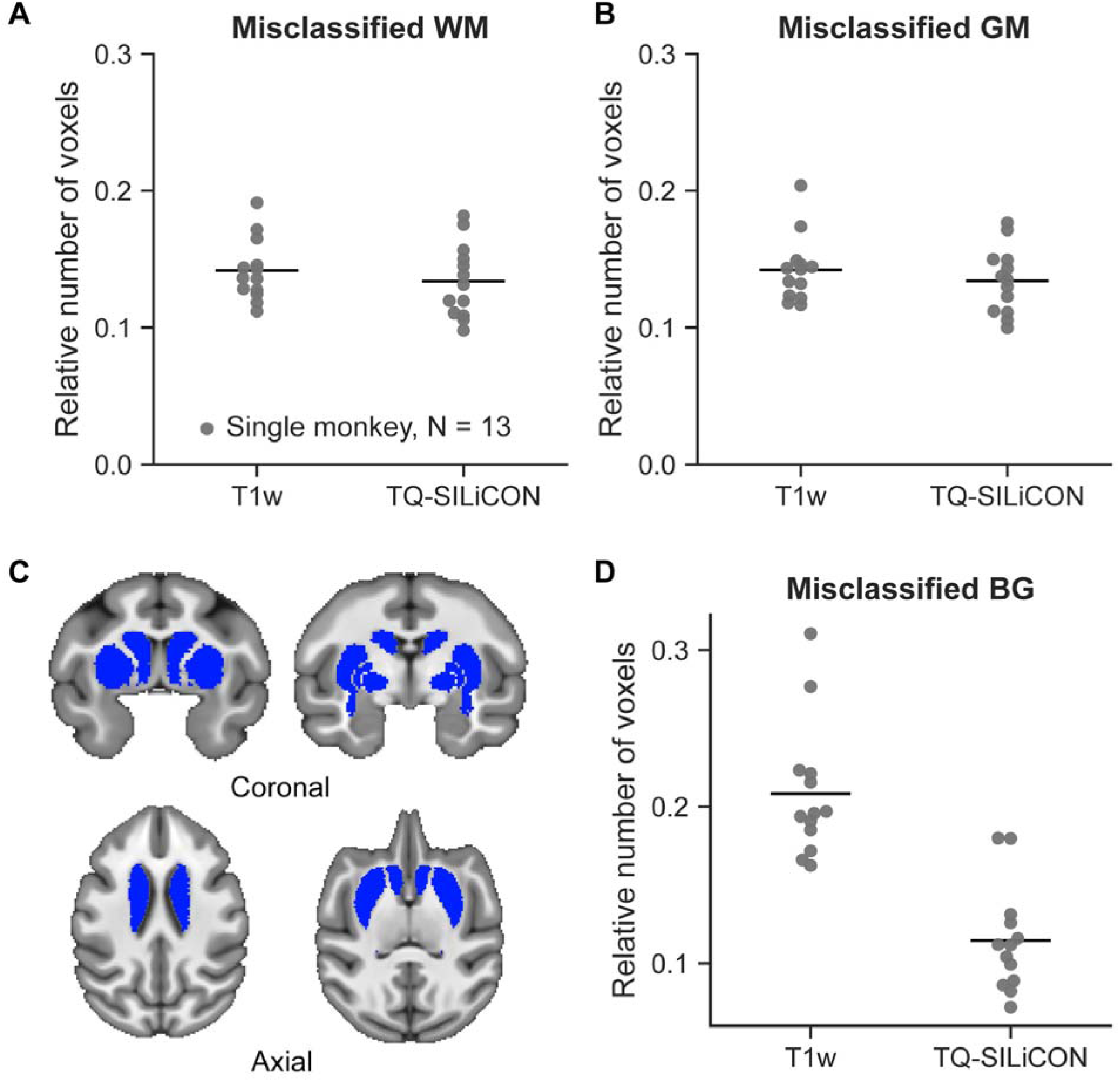
Improved single subject subcortical WM-GM classification. The total number of misclassified WM (**A**) and GM (**B**) voxels were normalized to the number of voxels WM and GM voxels available in the atlas, respectively. **C.** The basal ganglia (BG) mask was extracted from the D99 atlas and overlaid over the T1w average image. **D.** The total number of misclassified basal ganglia voxels were normalized to the total number of voxels in the basal ganglia mask. Whereas the number of WM voxels misclassification as GM was comparable between TQ-SILiCON and T1w. TQ-SILiCON produced significantly fewer misclassified voxels in the basal ganglia as compared to the T1w. GM - gray matter, WM - white matter, and BG – basal ganglia.

In summary, our fusion technique based on T1w and QSM resulted in a contrast-enhanced T1w-like 3D dataset that improved the automated segmentation algorithm and resulted in a more precise segmentation of gray and white matter in NHPs.

### 3.4 TQ-SILiCON enables the precise segmentation of human cortical and subcortical brain tissue

The TQ-SILiCON method also enabled an improved tissue segmentation of the human brain. By using the same weights as in macaques (e.g., 0.83 and −0.87), TQ-SILiCON of the human brain contained both excellent gray-white and deep gray matter contrast, as illustrated in **Fig. 7**. In comparison to the average T1w image, the average TQ-SILiCON revealed enhanced visibility and better delineation of subcortical structures. When comparing the average TQ-SILiCON to the average QSM map, the TQ-SILiCON image showed exceptional T1w-like gray-white contrast that the QSM image lacked (**Fig. 7**).

**Figure 7:**
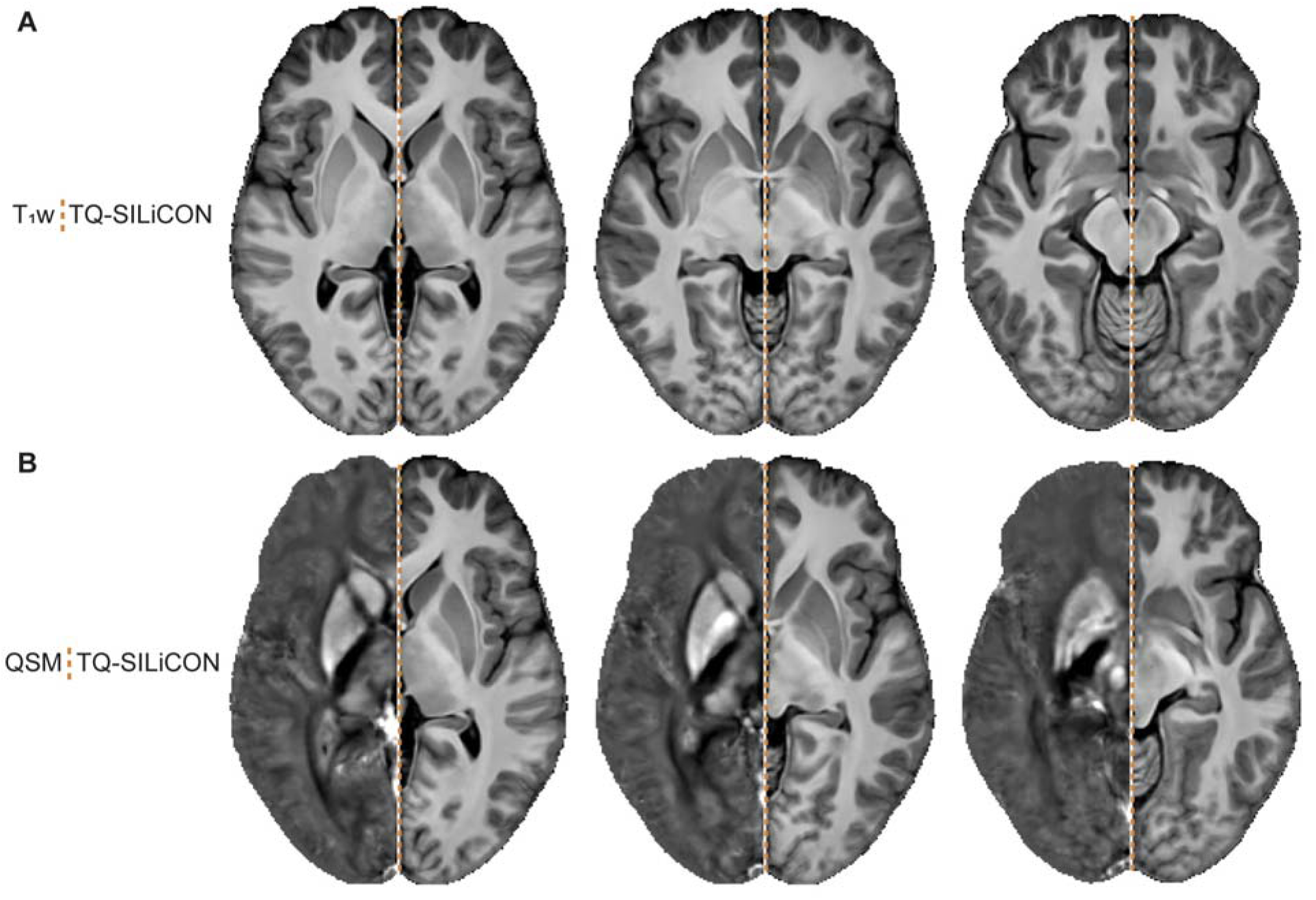
Enhanced cortical and subcortical contrast from TQ-SILiCON in the human brain **A.** Axially oriented TQ-SILiCON (right) slices of the average human template are compared side-to-side to the average T1w (left). **B.** Similarly, the TQ-SILiCON (right) is compared to QSM images (left). In comparison to T1w, TQ-SILiCON has enhanced subcortical contrast while maintaining high gray and white matter contrast, which the QSM map lacks.

The results of the single-subject segmentation in humans and the derived tissue probability masks are illustrated in **Fig. S3**. The TQ-SILiCON image-based segmentation correctly categorized the extreme capsule and external capsule as white matter and subcortical nuclei as gray matter, while the T1w image-based segmentation misclassifies gray matter nuclei as WM. In addition, T1w image-based segmentation misclassified white matter areas around the pre- and postcentral gyrus as gray matter in some human subjects, whereas TQ-SILiCON-based segmentation accurately classified gray matter structures from neighboring white matter areas (**Fig. S4**).

## 4. Discussion

In this study, we established an image contrast based on T1w and high-resolution QSM. This method, termed TQ-SILiCON, allowed us to automatically segment cortical and subcortical gray matter structures in NHPs and in humans. Given the reasonable measurement acquisition time, this contrast may facilitate translational studies in NHPs for investigating the pathophysiology of subcortical structures in humans. Below, we first discuss the advantages of QSM, including its potential for studying the basal-ganglia circuit of primates. We then discuss the advantages and limitations of combining QSM and T1w and how TQ-SILiCON could be used for neuroimaging applications in humans.

## Quantitative susceptibility mapping of the nonhuman primate brain

QSM is an MRI technique that relies on phase images to map the spatial distribution of paramagnetic and diamagnetic susceptibility (Haacke et al., 2005, 2004) that has just begun to be used in NHP neuroimaging (Yoshida et al., 2021). The enhanced contrast of brain tissue largely arises from different concentrations of iron, myelin, and calcium (Haacke et al., 2005). The absolute values derived from QSM could vary between MR systems and measurements as they could depend on the carrier frequency of the radio pulse, the B0 field, and the frequency drift (Liu et al., 2015). However, relative magnetic susceptibilities (e.g., relative to cerebrospinal fluid or the overall brain) have been shown to be reproducible and to be a common feature of QSM, particularly in the basal ganglia of humans (Li et al., 2011; Wharton and Bowtell, 2015). As described in healthy humans and during human development, QSM values have been linked to iron concentration (Bilgic et al., 2012). With increasing age, particularly subcortical deep gray matter structures show an increase in magnetic susceptibility in both humans (Bilgic et al., 2012; Keuken et al., 2017) and NHPs (Dadarwal et al., 2021a).

In contrast to humans, only a few studies have applied QSM in NHPs (Dadarwal et al., 2021a, 2019; Dadarwal and Boretius, 2021; Wen et al., 2020; Yoshida et al., 2021). The study by Yoshida et al. (Yoshida et al. 2021) reported an increased magnetic susceptibility in basal ganglia structures of the rhesus macaque, in accordance with our findings in the cynomolgus. Similarly, in our QSM maps, we observed high susceptibility in the caudate, putamen, globus pallidus (internal and external segments), thalamus, substantia nigra, red nucleus, and dentate nucleus of the cerebellum (**Fig. 3** and **4 A - B**). Importantly, while many of the enhanced subcortical structures are known to harbor iron, it is not well understood which substances and structures are most relevant for the bulk susceptibility. Identifying the sources that give rise to the increased susceptibility *in-vivo* has significant clinical implications for understanding the mechanisms that give rise to neurodegenerative diseases and aging (Ravanfar et al., 2021). Future immunohistochemistry studies in NHPs might provide a better understanding of the neural substrates and molecular mechanisms for the iron content and might enable means for quantifying the degree of iron contribution to the QSM map in the NHP brain.

Moreover, exploring cortical and subcortical structures with neuroimaging and interventional studies in NHPs (Klink et al., 2021) may improve our understanding of the functional properties of the cortico-striatal-thalamic circuits, further guiding therapeutic interventions. For example, the subthalamic nucleus (STN) is a typical target for deep brain stimulation (DBS) studies; its accurate delineation might aid in the physiological characterization of neuronal responses and microstimulation parameters; as has been shown in human patients undergoing DBS treatment (Liu et al., 2015). In addition, the delineation of the internal and external globus pallidus can aid in the MRI-guided targeting of the inhibitory and excitatory output of the cortico-striatal-thalamic circuit during micro-stimulation. The identification and visualization of small structures are likewise beneficial for neuroscientists implementing optogenetic (Galvan et al., 2017; Inoue et al., 2015) or chemogenetic interventions targeting deep brain nuclei (Mimura et al., 2021).

## Applications of TQ-SILiCON for brain segmentation

While QSM provides an outstanding contrast of deep gray nuclei, white matter structures are much better delineated on T1w. So, by linearly combining these two contrasts, we used “the best of two worlds” (**Fig. 4B**). As shown in the TQ-SILiCON image, our dual-fusion approach facilitated the delineation of the macaque brain’s cerebral cortex, white matter, and deep gray matter nuclei resulting in an overall T1w-like appearance. We believe TQ-SILiCON can aid stereotactic interventions of small subcortical structures such as the subthalamic nucleus and can aid in the delineation of the subcortical deep gray matter nuclei while maintaining adequate anatomical information of the whole brain is crucial for precise targeting.

Another useful application for the TQ-SILiCON is for the development of brain atlases in humans and NHPs. T1w is typically used in human and NHP brain atlases due to its excellent white and gray matter contrast. However, given that T1w alone is insufficient, neuroimaging studies in humans explored the use of additional MRI contrasts such as T2, T2*, and QSM to generate contrast from subcortical deep gray matter nuclei (Alkemade et al., 2017; Bazin et al., 2020; Xiao et al., 2015). The available macaque atlases such as the D99 (Saleem et al., 2021; Saleem and Logothetis, 2012) and the NMT (Hartig et al., 2021; Jung et al., 2021) incorporate parcellations based on T1w only and cytoarchitectonic mapping derived from post-mortem histology. In the future, we believe that adding multi-contrast acquisitions might aid in the development of more precise atlas parcellations and clearer delineation of regional boundaries, although with the limitation of an increase in acquisition time. However, our proposed TQ-SILiCON approach might be a more viable option for obtaining a shorter in-session acquisition.

The accurate segmentation of gray and white matter tissue is not only essential for basic neuroscience applications but also for clinical diagnostics. For example, during brain development, aging, and neurological diseases, specific volumetric changes might occur that might require precise segmentation and delineation of the overall structure (Boedhoe et al., 2020; Liu et al., 2015). Importantly, algorithms for automatic brain tissue segmentation typically utilize template-derived tissue probability masks (CSF, gray matter, and white matter) to initiate the segmentation process of a single subject volume. TQ-SILiCON atlases may aid in the generation of more accurate probability masks. As demonstrated by our automated tissue segmentation analyses. In our study, TQ-SILiCON outperformed T1w based tissue segmentation in both macaques (**Fig. 5)** and humans (**Fig. S3**). Interestingly, TQ-SILiCON not only provided a better delineation of subcortical gray matter structures but also improved the assignment of white matter structures such as the extreme and external capsule and white matter areas around the pre-and postcentral gyrus (**Fig. S2** and **S4**), even in single subjects and across different subject ages. However, one limitation of our TQ-SILiCON approach relates to the segmentation of the CSF areas, which occasionally exhibited a comparable contrast to the basal ganglia. In our analyses, we bypassed this limitation by segmenting the CSF based on T1w and then extracting the volume from TQ-SILiCON before applying the segmentation process. In the future, the TQ-SILiCON approach could be improved by incorporating more than two contrasts to attain higher segmentation accuracy. An advantage of the TQ-SILiCON, particularly for its application in humans as opposed to the advanced deep learning based multicontrast MR image synthesis approach (Yu et al., 2020), is the simplicity of implementation, which does not require extensive computational resources. The linear combination of T1w and QSM contrasts preserves the original data features, which are retained in the TQ-SILiCON image.

In summary, we believe that TQ-SILiCON can significantly improve the delineation of brain structures and the accuracy of morphometric studies. T1w and QSM data sets could be obtained using clinically available 3T-MR-systems and in a reasonably short acquisition time. Importantly, the method works equally well for NHPs and humans, facilitating translational studies. In the future, we will aim to study the structural underpinnings of the MRI contrast observed *in-vivo* and hope to contribute to a more comprehensive understanding of the underlying pathophysiology behind neurodegenerative diseases.

## 6. Supplementary Materials

**Figure S1:**
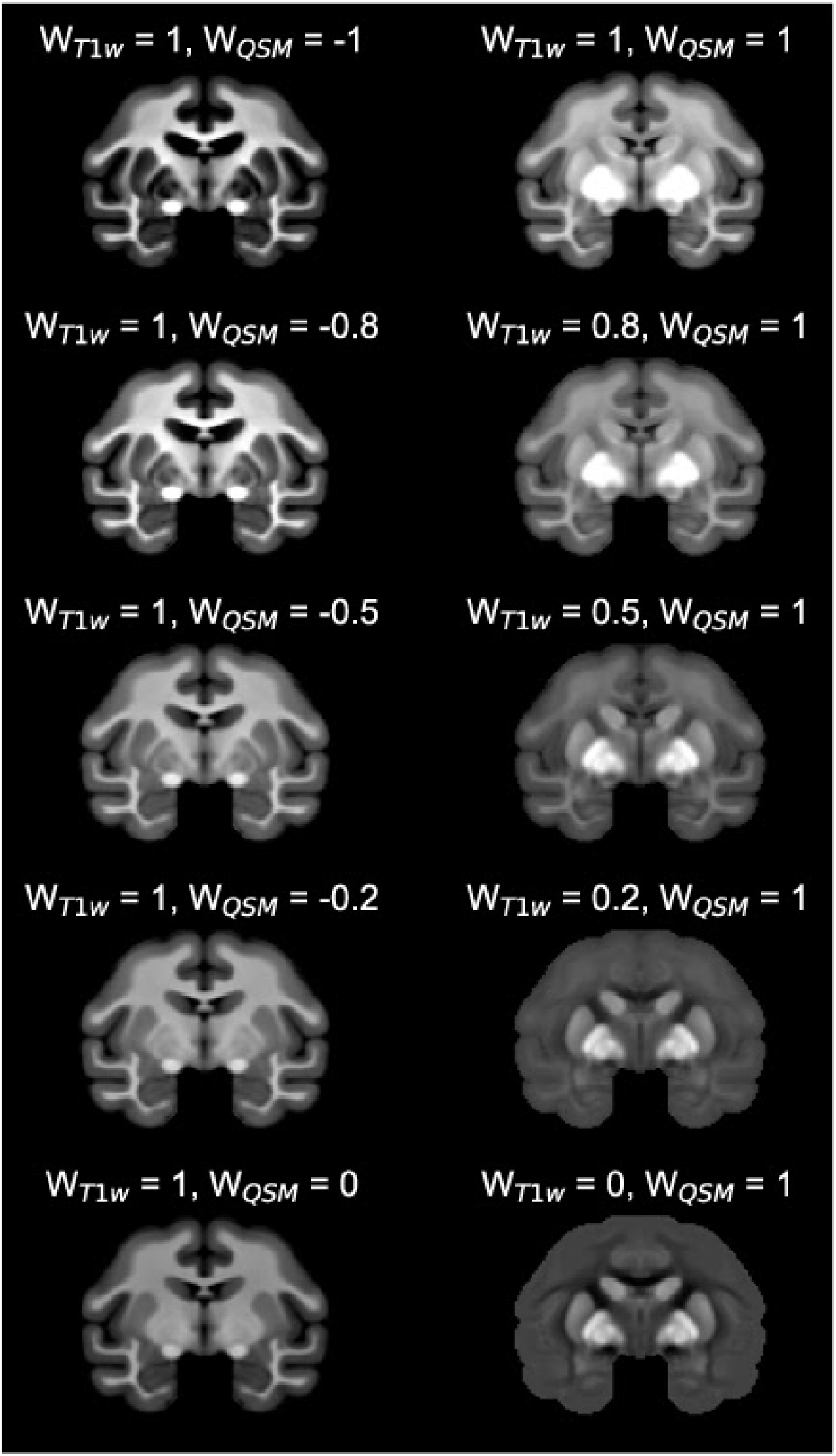
Examples of differently weighted linear combinations of T1w and QSM.

**Figure S2:**
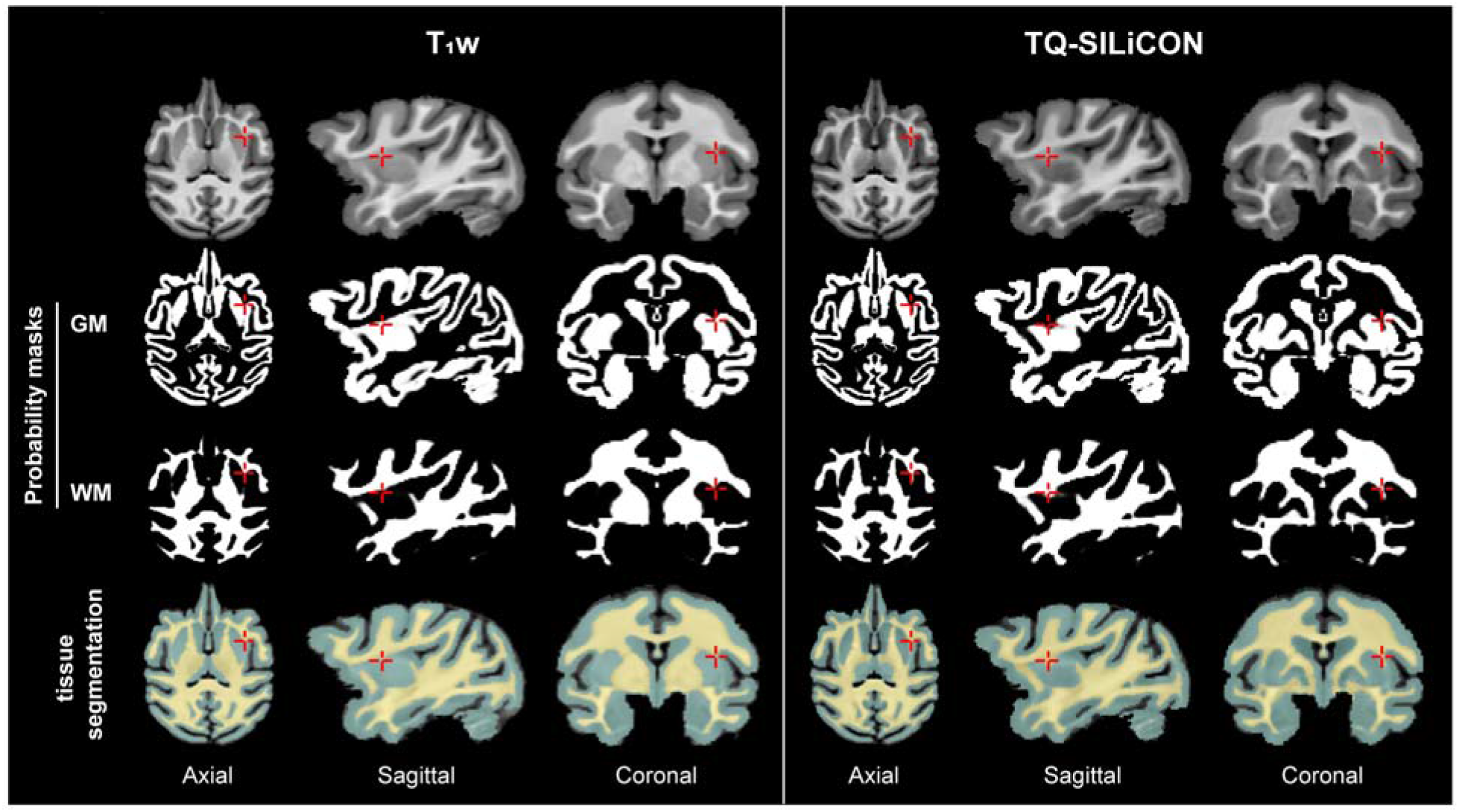
T1w and TQ-SILiCON image segmentation results for a single macaque brain. T1w segmentation misclassified some white matter areas as gray matter (highlighted area in the external capsule). TQ-SILiCON image segmentation, on the other hand, improved cortical gray/white matter delineation. Abbreviations: GM - gray matter, and WM - white matter.

**Figure S3:**
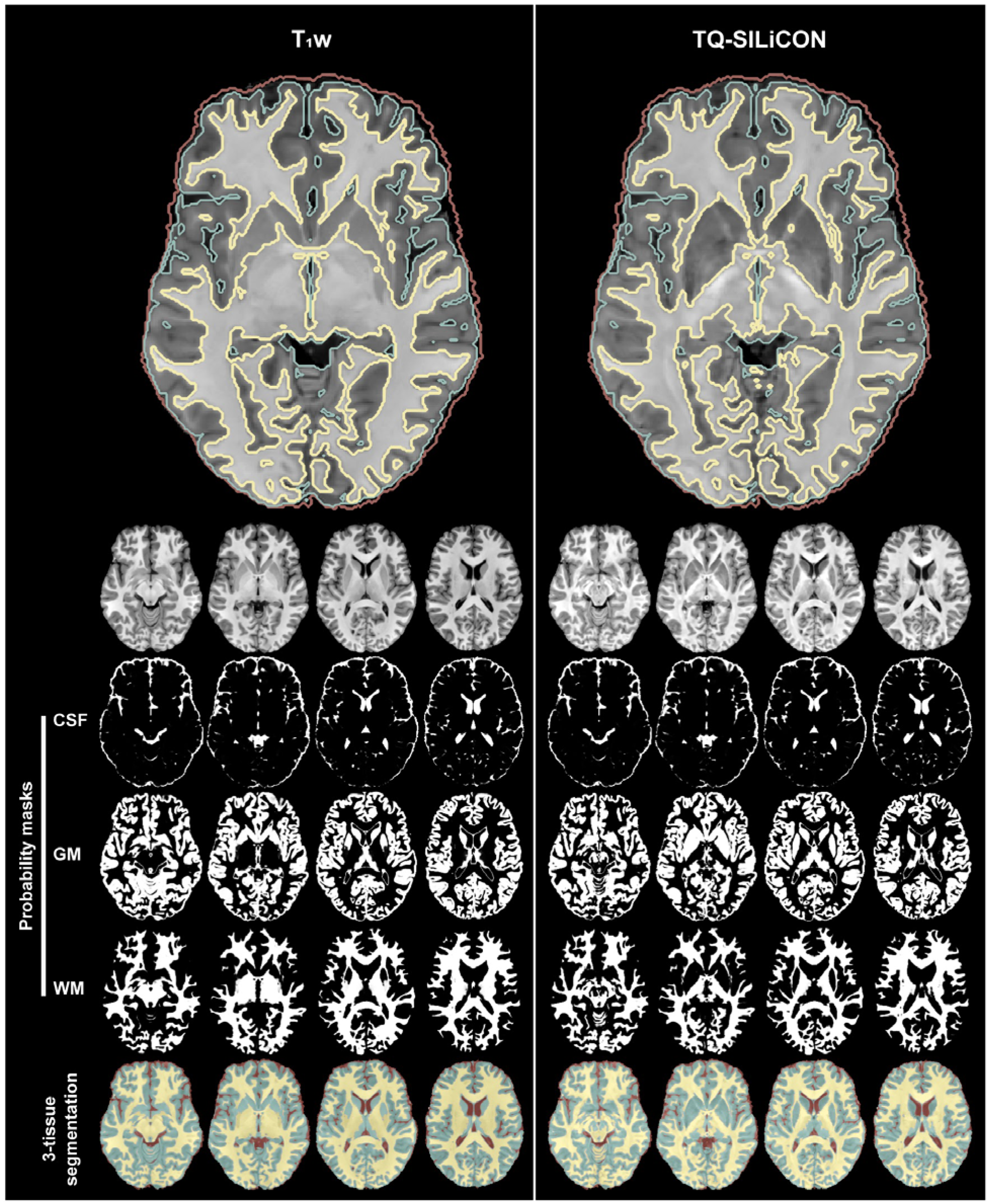
Axial view of single human brain tissue segmentation using T1w (left) and TQ-SILiCON images (right). TQ-SILiCON enables better tissue classification of gray matter and white matter. The bottom shows the semi-transparent color-coded tissue classification of the CSF, WM, and GM. Abbreviations: CSF - cerebrospinal fluid, GM - gray matter, and WM - white matter.

**Figure S4:**
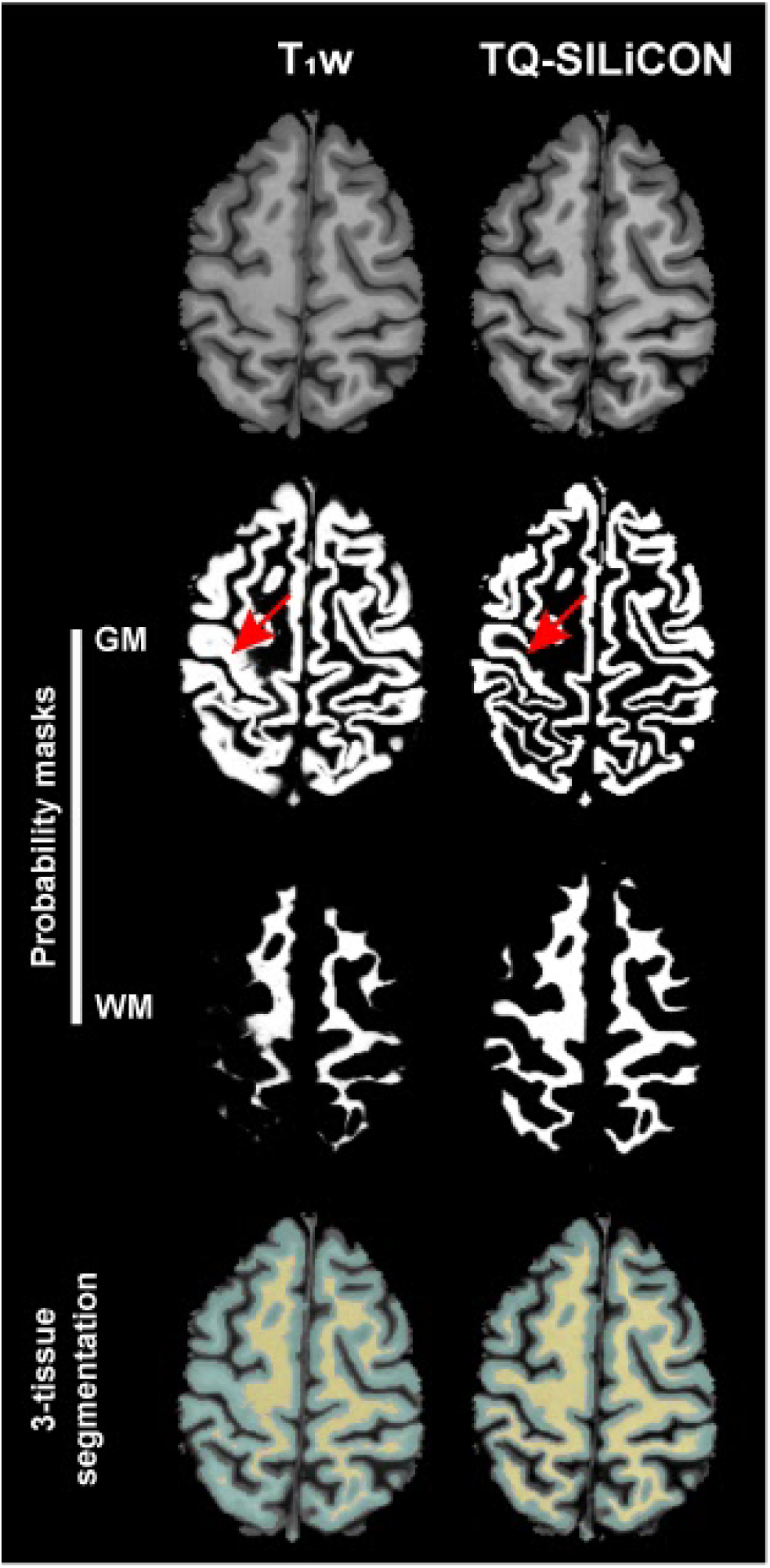
The axial view of T1w and TQ-SILiCON images of a single human brain and associated GM and WM probability masks and segmentation images. Compared to the T1w image, TQ-SILiCON imagebased segmentation better delineates gray and white matter. Abbreviations: GM - gray matter, and WM - white matter.

**Figure S5:**
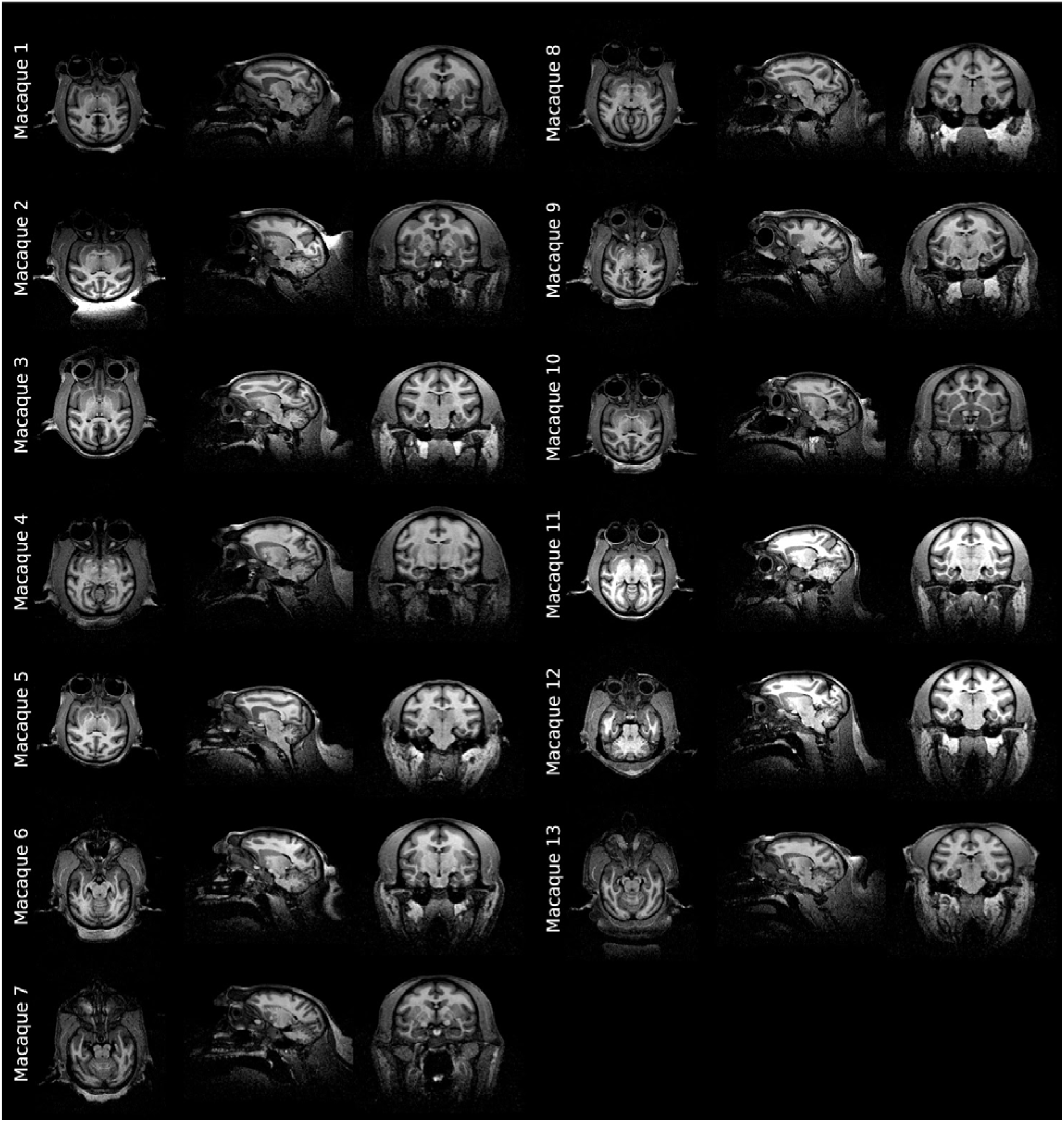
Axial, sagittal, and coronal views of the raw T1w images for the 13 macaques used in the study. All data were obtained under anesthesia (movement-related artifacts) and in naive individuals (e.g., non-implanted), resulting in artifact-free imaging data.

**Figure S6:**
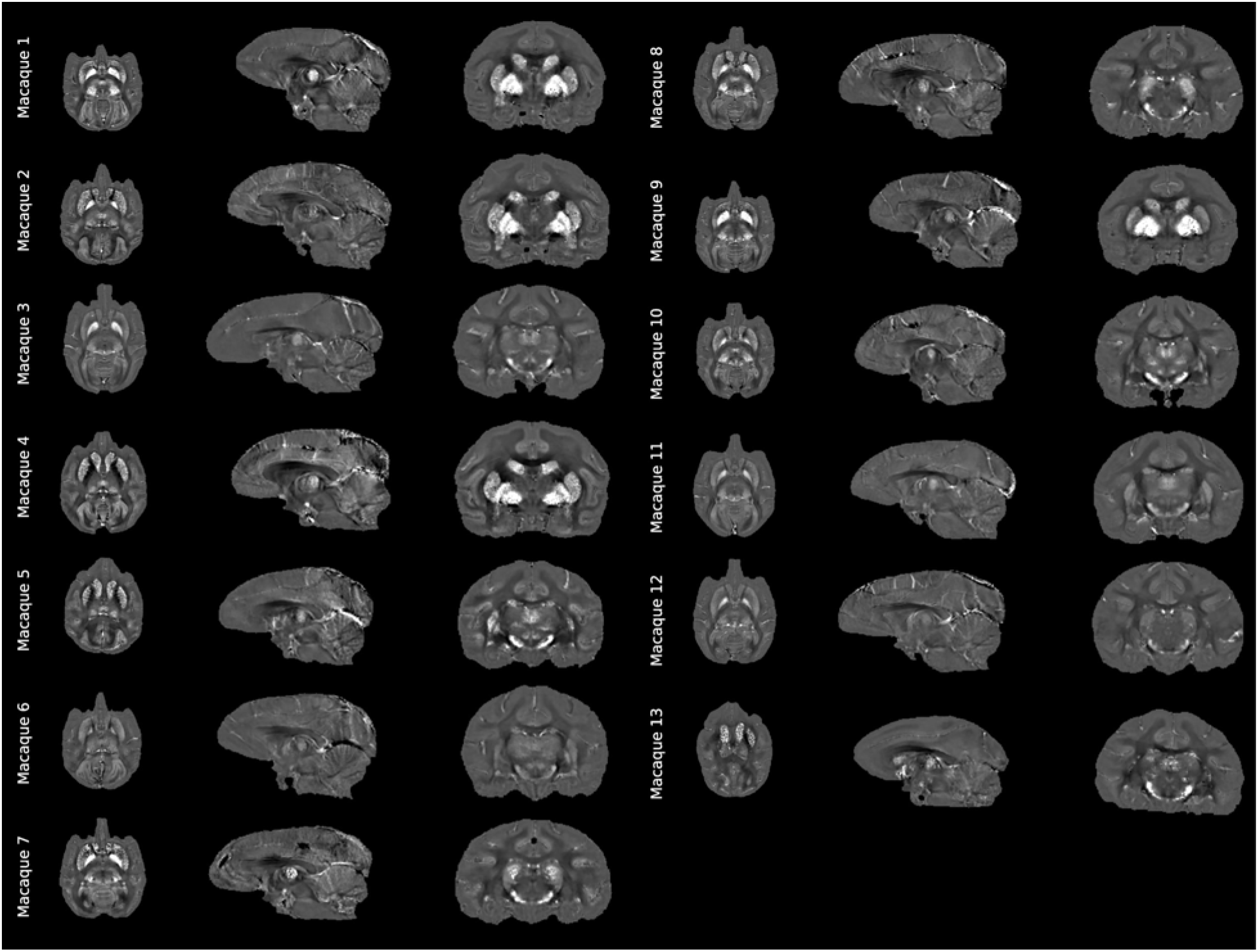
Axial, sagittal, and coronal views of the QSM maps for the 13 macaques used in the study.

**Figure S7:**
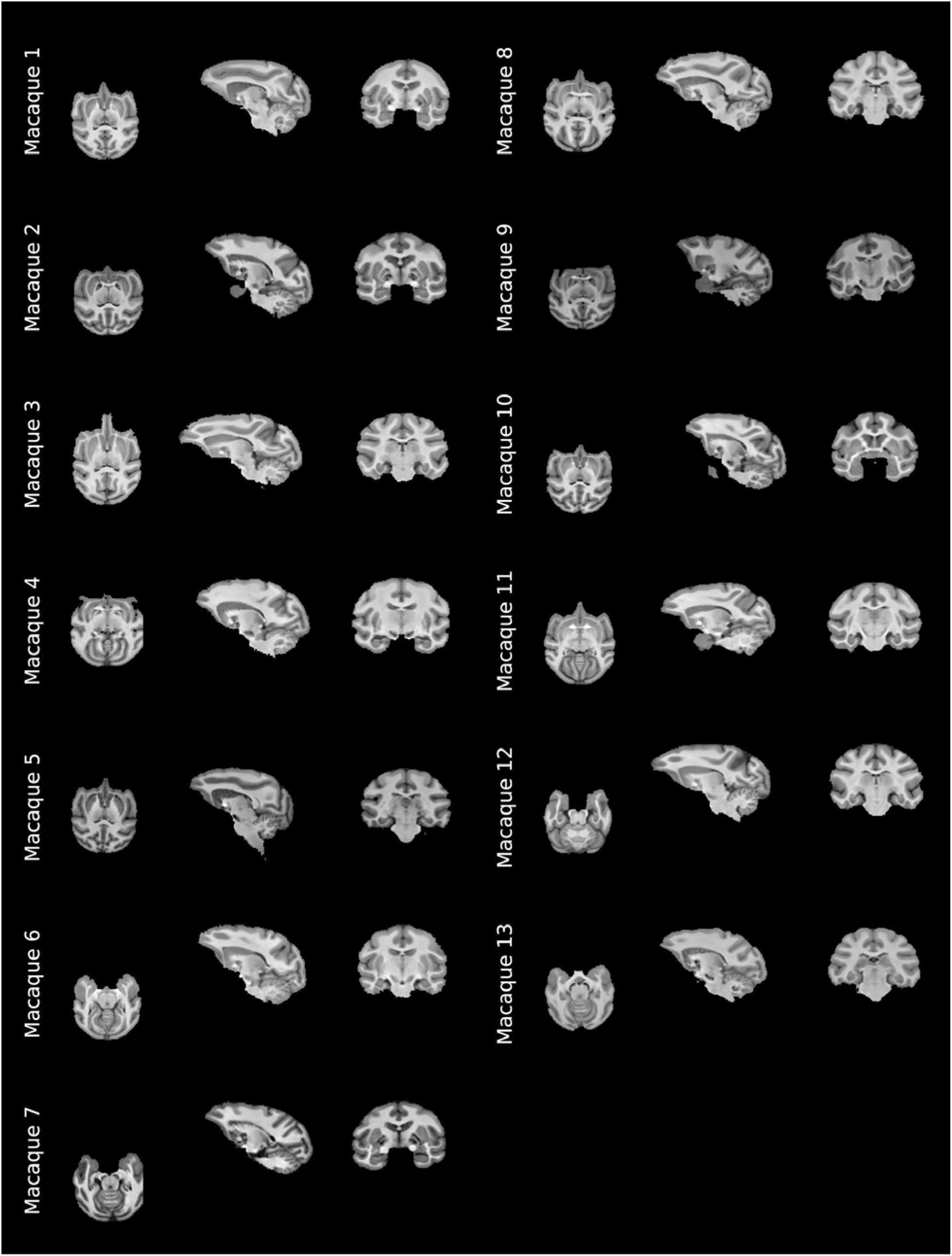
Axial, sagittal, and coronal views of the TQ-SILiCON maps for the 13 macaques used in the study.

## Notes

### Competing Interest Statement

The authors have declared no competing interest.

